# Dendritic membrane resistance modulates activity-induced Ca^2+^ influx in oxytocinergic magnocellular neurons of mouse PVN

**DOI:** 10.1101/2021.01.26.428216

**Authors:** Wanhui Sheng, Scott W. Harden, Yalun Tan, Eric G. Krause, Charles J. Frazier

**Affiliations:** Department of Pharmacodynamics, College of Pharmacy, University of Florida; Department of Anesthesiology, School of Medicine, Stanford University; Center for Integrative Cardiovascular and Metabolic Diseases, University of Florida, Gainesville, FL; Evelyn F. and William L. McKnight Brain Institute, University of Florida, Gainesville, FL

**Keywords:** oxytocin, hypothalamus, magnocellular neuron, PVN, dendrite, osmotic stress, electrophysiology

## Abstract

Hypothalamic oxytocinergic magnocellular neurons have a fascinating ability to release peptide from both their axon terminals and from their dendrites. Existing data indicates there is a flexible relationship between somatic activity and dendritic release, but the mechanisms governing this relationship are not completely understood. Here we use a combination of electrical and optical recording techniques to quantify activity-dependent calcium influx in proximal vs. distal dendrites of oxytocinergic magnocellular neurons located in the paraventricular nucleus of the hypothalamus (OT-MCNs). Results reveal that the dendrites of OT-MCNs are weak conductors of somatic voltage changes, and yet activity-induced dendritic calcium influx can be robustly regulated by a diverse set of stimuli that open or close ionophores located along the dendritic membrane. Overall, this study reveals that dendritic membrane resistance is a dynamic and endogenously regulated feature of OT-MCNs that is likely to have substantial functional impact on central oxytocin release.

**IMPACT STATEMENT:** Activity-induced dendritic calcium influx in oxytocinergic magnocellular neurons can be robustly modulated by a highly diverse set of stimuli acting on distinct types of ionophores expressed along the dendritic membrane.

## INTRODUCTION

Oxytocin (OT) is a nine amino acid peptide synthesized almost exclusively in hypothalamic neurons of the supraoptic and paraventricular nucleus (SON, PVN). Despite the highly localized nature of OT synthesizing neurons, OT receptors (OTRs) are distributed widely throughout the CNS. Significant evidence indicates an important modulatory role for central oxytocinergic signaling in a wide variety of processes including fear conditioning, stress responding, anxiety-related behaviors, maternal behavior, sexual behavior, and social recognition (e.g. see Bosch et al., 2005; Knobloch et al., 2012; Jurek et al., 2015; Veening et al., 2015; Neumann and Slattery, 2016; Lin et al., 2018; Tan et al., 2019; Valtcheva and Froemke, 2019; Winter and Jurek, 2019). Numerous studies have demonstrated notable effects of OTR agonists and antagonists delivered intracerebrally or intraventricularly on social, stress, and anxiety-related behaviors, while conversely, animal models with genetic disruptions in OT signaling systems exhibit a range of social and behavioral deficits (e.g. see Jin et al., 2007; Young, 2007; Lee et al., 2008; Higashida et al., 2010; Pobbe et al., 2012; Kent et al., 2013; Morales-Rivera et al., 2014; Peters et al., 2014; Burkett et al., 2016; Caldwell et al., 2017; Lee et al., 2018). Collectively, these types of data implicate the central OT signaling system as a promising therapeutic target for a variety of conditions impacting mental health. However, effective therapeutic delivery of exogenous OTR agonists into the CNS of humans remains difficult (Ermisch et al., 1985; McEwen, 2004; Veening and Olivier, 2013; Leng and Ludwig, 2016; Quintana and Woolley, 2016; De Cagna et al., 2019), and thus a more detailed mechanistic understanding of how the brain naturally regulates release of endogenous OT may facilitate development of new approaches for therapeutic manipulation of central OT signaling.

The question of exactly how, and from where, endogenous OT is released to act on both hypothalamic and extrahypothalamic OTRs in the CNS is a complex one. It is complicated by the fact that there are both magnocellular and parvocellular hypothalamic OT synthesizing neurons (OT-MCNs, OT-PCNs), and by the fact that OT-MCNs can release peptide both from their axon terminals and from their dendrites. The current study focuses on dendritic physiology of PVN OT-MCNs because 1) OT-MCNs substantially outnumber OT-PCNs (Althammer and Grinevich, 2017), 2) most axons of OT-MCNs project through the median eminence to the posterior pituitary where activity (action potential) dependent release into the vasculature increases peripheral (and not central) OT concentration (Guzek, 1987; Robinson et al., 1989; Falke, 1991), 3) a large portion of OT available for release into the CNS clearly exists in dendritic rather than axonal vesicles that are subject to activity, and calcium, dependent exocytosis (Pow and Morris, 1989; Ludwig et al., 2002; Ludwig and Leng, 2006), 4) such exocytosis supports functionally important peptide mediated paracrine signaling within the hypothalamus (Son et al., 2013; Smith et al., 2015; Pati et al., 2020), and 5) likely also drives volume transmission to increase functional activation of OTRs in a variety of extrahypothalamic cortical and limbic areas (Veening et al., 2010; Fuxe et al., 2012; Ludwig and Stern, 2015; Brown et al., 2020). Indeed, this mechanism seems likely to work in concert with limited/targeted release from centrally projecting axon collaterals of OT neurons, as has been effectively demonstrated in several extrahypothalamic areas to date (Knobloch et al., 2012; Eliava et al., 2016; Oettl et al., 2016).

In the current study we use a combination of electrophysiological and subcellular optical recording techniques to evaluate dendritic physiology of OT-MCNs. Somatic activity was simulated with a reproducible train of action potential-like voltage pulses delivered to the soma, and calcium influx induced by this somatic activity was quantified using high frequency two-photon line scans across the proximal and distal dendrite. The results reveal that the dendrites of OT-MCNs are weak conductors of somatic voltage changes. We further report that hyperosmotic stress preferentially reduces activity-dependent calcium influx in distal vs. proximal OT-MCN dendrites, while hypoosmotic stress increases it. Extensive control experiments indicate these effects are likely mediated by modulation of a previously unidentified osmosensitive channel expressed along the dendritic membrane. Finally, we report that that activity-induced calcium influx in the distal dendrites of OT-MCNs is also preferentially and robustly inhibited, absent any change in osmotic stress, by activation of dendritic GABA_A_ receptors. Collectively, these results significantly increase our understanding of the mechanisms through which the brain is likely to dynamically regulate the relationship between somatic activity and calcium-dependent dendritic release of OT into the CNS.

## MATERIALS AND METHODS

### Animals

All experiments in this study were performed using 1-3-month-old OT-reporter mice which expresses red fluorescent protein (tdTomato) in oxytocinergic neurons. These mice were generated by crossing OT-IRES-Cre knock-in mice (Jackson Labs Stock #024234) with Ai14 mice that express tdTomato following a loxP-flanked stop cassette in the Rosa26 locus under control of the CAG promoter (Jackson Labs Stock #007914). This same strategy has been used in several previous studies to facilitate identification of oxytocinergic neurons (Clipperton-Allen et al., 2016; Xiao et al., 2017). Throughout the course of this study experiments were performed on approximately equal numbers of male and virgin female mice. Results of all core experiments, including immunohistochemical evaluation of OT-reporter animals, analysis of intrinsic properties of OT-MCNs, and the evaluation of the effects of acute osmotic stress on OT-MCNs, observed both somatically and in the distal dendrites, were carefully compared across sex. Because we noted no major sex-based differences between males and virgin females in any of these experiments, data in all primary figures are combined across sex. See main text of results section and Extended Data Figures 1-1, 2-1, and 4-1 for further details. All animals were group-housed on a 12 hr light/dark cycle, and all animal procedures were approved by the University of Florida Institutional Animal Care and Use Committee (IACUC).

### Immunohistochemistry

We used immunohistochemical techniques to quantify co-expression of tdTomato and oxytocin-neurophysin 1 (NP1). Two male and two female mice were anesthetized with sodium pentobarbital (1.56 mg/g i.p.) and transcardially perfused with 0.15 M NaCl followed by 4% paraformaldehyde. Brains were extracted post-fixed for 4 hours in 4% paraformaldehyde, then transferred to a sucrose solution (30% sucrose in PBS) and stored at 4 °C for at least 24 hr. Brains were then sectioned at 30 microns and stored at −20 °C in cryoprotective solution (1 L of 0.1 M PBS supplemented with 20 g PVP-40, 600 mL ethylene glycol, and 600 g sucrose). Unless otherwise noted, immunohistochemistry was conducted at room temperature using free-floating sections in 12-well plates (3 mL / well) on an orbital shaker. Following 5 rinses (5 min each) with 50 mM potassium PBS (KPBS), tissue was incubated in blocking solution (KPBS with 2% normal donkey serum and 0.2% Triton X-100) for 1 hour. Subsequently, sections were incubated in a primary antibody against NP1 overnight at 4 °C (mouse monoclonal, PS-38, Dr. H. Gainer, National Institute of Health, 1:400)(Ben-Barak et al., 1985; de Kloet et al., 2016). Following 5 rinses (5 min each) in KPBS, sections were incubated in the secondary antibody (Alexa 647 Donkey anti-mouse, Jackson Immuno 715-605-150, 1:500) for 2 hours. Following 5 rinses (5 min each) with KPBS, slices were mounted on glass slides, air dried overnight, and coverslipped with polyvinyl alcohol mounting medium. Images were captured and analyzed using a Nikon C2+ scanning confocal microscope and NIS-Elements AR 5.02 software. Identical imaging parameters were used for the acquisition and analysis all images.

### Acute Brain Slice Preparation

Mice received an IP injection of ketamine (0.1 mL of 100 mg / mL, in sterile physiological saline) and were euthanized using a small animal guillotine. Brains were extracted and coronal sections 200 μm thick were made using a Leica VT1000 S vibratome. During this procedure, slices were submerged in ice-cold, sucrose-laden, artificial cerebrospinal fluid (ACSF) containing (in mM): 87 NaCl, 2.5 KCl, 1.25 NaH_2_PO_4_, 7 MgCl_2_, 10 dextrose, 0.5 CaCl_2_, 75 sucrose, and 25 NaHCO_3_. After sectioning was complete, brain slices containing the PVN were transferred to an incubator filled with a low-calcium high-magnesium ACSF that contained (in mM): 124 NaCl, 2.5 KCl, 1.23 NaH_2_PO_4_, 10 dextrose, 1 CaCl_2_, 3 MgSO_4_, and 25 NaHCO_3_. Both solutions were continuously saturated with 95% O_2_/5% CO_2_ and had a pH of ~7.3. After 30 minutes of incubation at 37 °C, slices were passively equilibrated to room temperature for an additional 30 minutes (minimum) prior to use.

### In Vitro Electrophysiology

In preparation for in vitro electrophysiological experiments, slices were transferred from the slice incubator to a low turbulence perfusion chamber (JG-23W/HP, Warner Instruments) where they were continuously perfused a rate of 2 mL/min with ACSF containing (in mM) 126 NaCl, 11 dextrose, 1.5 MgSO_4_, 3 KCl, 1.2 NaH_2_PO_4_, 2.4 mM CaCl_2_, and 25 NaHCO_3_. This solution was continuously oxygenated with 95% O_2_ and 5% CO2, had a pH of ~7.3, and was maintained at 28 °C. tdTomato positive PVN neurons were identified using an Olympus BX51WI stereomicroscope that supported both infrared differential interference contrast (IR-DIC) and conventional epifluorescence microscopy. Both IR-DIC and epifluorescence images were acquired through an Olympus 40X water immersion objective using 12-bit IR CCD camera (QICAM Fast 1394) controlled by Fiji software (Schindelin et al., 2012). An X-Cite Series 120Q (Lumen Dynamics) light source coupled with an XF406 filter set (Omega Optical) was used for conventional epifluorescence imaging. Patch pipettes were prepared using a Flaming/Brown pipette puller (Sutter Instruments, P-97). Borosilicate glass capillaries (1.5 mm/0.8 mm) were pulled to produce patch pipettes with an open tip resistance of 4-6 MΩ when filled with an intracellular solution that contained (in mM): 10 KCl, 5 NaCl, 2 MgCl_2_, 1 EGTA, 10 HEPES, 130 K-gluconate, 10 phosphocreatine, 2 Na2-ATP, 0.3 Na-GTP, adjusted to pH 7.25 and 290 mOsm. This solution was used for all experiments that involved whole-cell recording but not simultaneous calcium imaging. For experiments that required concurrent calcium imaging (see next section), EGTA was omitted and replaced with 0.3 mM Fluo-5F (ThermoFisher F14221) and 0.03 mM Alexa 594 (ThermoFisher A10438). Whole-cell recordings were made using a Multiclamp 700B amplifier, Digidata 1440A digitizer, and Clampex 10.7 software (Molecular Devices). All whole-cell data were sampled at 20 kHz and low-pass filtered at 2 kHz. When necessary, synaptic responses were generated using a small tipped bipolar stimulator pulled from theta glass (~1-1.5 μm inner diameter), connected to a constant current stimulus isolator (0.2 ms pulse duration). Simulator placement and stimulation intensity were adjusted until an evoked response ≥ 50 pA could be reliably generated. Online analysis of whole-cell patch clamp data was performed with custom software using Python 3.6 and the pyABF module. Off-line analysis was performed using custom software written by CJF in OriginC (OriginLab Corporation, Northampton, MA). All experiments were performed in the continuous presence of bath applied antagonists for kainate/AMPA receptors, NMDA receptors, GABA_A_ receptors, and GABA_B_ receptors (20 μM DNQX, 40 μM AP5, 100 μM picrotoxin (PTX), and 10 μM CGP-55845 (CGP), respectively), except for those that explicitly involved measuring evoked excitatory synaptic currents (EPSCs), or effects of an exogenous glutamate or GABA receptor agonist. Some experiments transiently delivered tetrodotoxin (TTX, 1 μM), muscimol (400 nM), or ruthenium red (RR, 10 μM) using a syringe pump in-line with the bath perfusion system. In experiments that involved changing osmolarity of the bath, mannitol (MT, in ACSF) or purified deionized water were also delivered using a syringe pump. Local application of glutamate was achieved using a Parker Picospritzer III (model R374-01C, 5-10 ms pulse duration, 10-20 psi), connected to a glass pipette, identical to those used for whole-cell recording, loaded with 100 mM glutamate solubilized in ACSF. All drugs used during in vitro experiments were obtained from TOCRIS except PTX and glutamate which were obtained from Sigma-Aldrich. Although stock solutions for PTX and CGP were dissolved in DMSO, total bath concentration of DMSO never exceeded ~0.1% and remained stable throughout experiments.

### Subcellular calcium imaging

Subcellular two-photon calcium imaging in OT-MCN axons and dendrites was accomplished using an Ultima laser scanner (Bruker Scientific, Billerica, MA), using a Coherent Mira Ultrafast Ti:sapphire laser (powered by a 5W Coherent Verdi laser, Coherent, Inc. Santa Clara, CA). The emission wavelength of the Mira was set to 810 nm in order to excite both the calcium sensitive indicator, Fluo-5F, and the calcium insensitive indicator, Alexa Fluor 594 (see above for detailed composition of internal solution). All cells were permitted to rest for 15 minutes following establishment of whole-cell configuration to ensure robust diffusion of fluorophore into dendrites before proceeding with calcium imaging experiments. Somatic activity was induced in cells voltage-clamped at −70 mV using a train of brief action potential like voltage steps (to +50 mV for 5 ms, delivered at 20 Hz for 2 seconds). Activity-dependent calcium influx was measured in dendrites or axons using two-photon line scans across specific structures of interest (11.9 s duration, 85 Hz) at known distances from the soma. Emissions from Fluo-5F (green, G) and Alexa Fluor 594 (red, R) were separated with a dichroic mirror and simultaneously measured by two separate photomultiplier tubes (PMTs, Hamamatsu R3896 SEL). Data from the PMTs was used to calculate G/R, which was then baseline subtracted to the pre-stimulus baseline, smoothed with a Gaussian window function, and reported as ΔG/R. Cells were discarded if pre-stimulus calcium increased by ≥ 20% during the course of an experiment. Dendrites were distinguished from axons by their clearly larger diameter and by their enhanced passive fill with somatically delivered Alexa Fluor 594. Off-line analysis of line scan data was performed using custom software written in C# and OriginC by SWH and CJF. Where mentioned in the text, the voltage clamp current applied to the soma during the train was quantified over time by measuring the area under the curve of the last pulse.

### Statistical Analysis

Within-cell changes in individual parameters produced by an experimental procedure were evaluated with a two-tailed 1-sample Student’s t-test using data normalized to the baseline mean (null hypothesis: test mean=1). When comparing a parameter of interest across two independent groups of cells a two-tailed 2-sample Student’s t-Test was used (null hypothesis: Group 1 mean = Group 2 mean). Welch’s correction was applied in cases where population variance was significantly different between samples. For comparisons that involved multiple measurements per cell (e.g. delay to first spike as a function of injected current, or activity-induced ΔG/R observed in the dendrite at > 2 distances) a one-way or two-way repeated measures ANOVA was employed as necessary, with Holm-Sidak post-hoc tests where appropriate. Extent of co-localization of tdTomato and NP1 in males vs. females was evaluated using a two-sample proportion test. In all cases p-values ≤ alpha level of 0.05 were considered statistically significant. Error bars on all plots represent the SEM.

## RESULTS

### Identification of OT-MCNs and PCNs in vitro

All experiments for this study were performed in an OT-reporter mouse line designed to selectively expresses a red fluorescent protein (tdTomato) in oxytocinergic neurons (Figure 1A, see Methods). In order to validate specificity and selectivity of this reporter in the PVN, we used immunohistochemical techniques to evaluate co-expression of tdTomato and neurophysin 1 (NP1, an OT carrier protein found in oxytocinergic neurons, Figure 1B). Across all animals tested (n=2 male, 2 female), we found that over 92% of tdTomato-expressing neurons in PVN were immunoreactive for NP1 (2629/2838). Importantly, we also noted that 1262/1352 and 1367/1468 of tdTomato positive neurons examined were NP1 positive in males and females, respectively, indicating that there is no sex based difference in the extent of co-localization (Z=1.38, p=0.17). Based on these data, we used a combination of IR-DIC and epifluorescence microscopy (see Fig. 1C and Methods) to efficiently target PVN OT neurons for whole-cell patch clamp recording. Once patched, PVN OT neurons were categorized as either magnocellular or parvocellular based on the presence or absence (respectively) of a transient outwardly rectifying potassium current, I_A_ (Luther and Tasker, 2000). This current was readily apparent as an outward current in voltage clamp observed in the first 100 ms after stepping from −70 mV to −50 mV (Fig. 1D-E). It also contributed to a clear delay to first action potential observed in current clamp in response to a suprathreshold current injection (Fig. 1F-G). By contrast, OT PCNs not only lacked I_A_, but often also displayed a small inward current after stepping from −70 mV to −50 mV in voltage clamp (Fig. 1D, bottom trace). This current often produced a low threshold spike that was apparent in current clamp (Fig. 1F, bottom trace), and contributed to the short delay to first spike observed in OT PCNs (Fig. 1G). We noted that OT-MCNs had significantly higher somatic input resistance than OT-PCNs (1836.87 ± 107.84 MΩ vs. 771.9 ± 81.7 MΩ, respectively, n=62, 16,, t=7.9, p<0.0001), while whole-cell capacitance at −70 mV did not significantly differ (Cm: 31.1 ± 1.4 pF vs. 31.3 ± 2.3 pF, n=62, 16, t=-0.07, p=0.94). Importantly, none of the core intrinsic electrophysiological features of OT-MCNs reported here varied by sex (See Extended Data Fig. 1-1 and legend).

**Figure 1.**
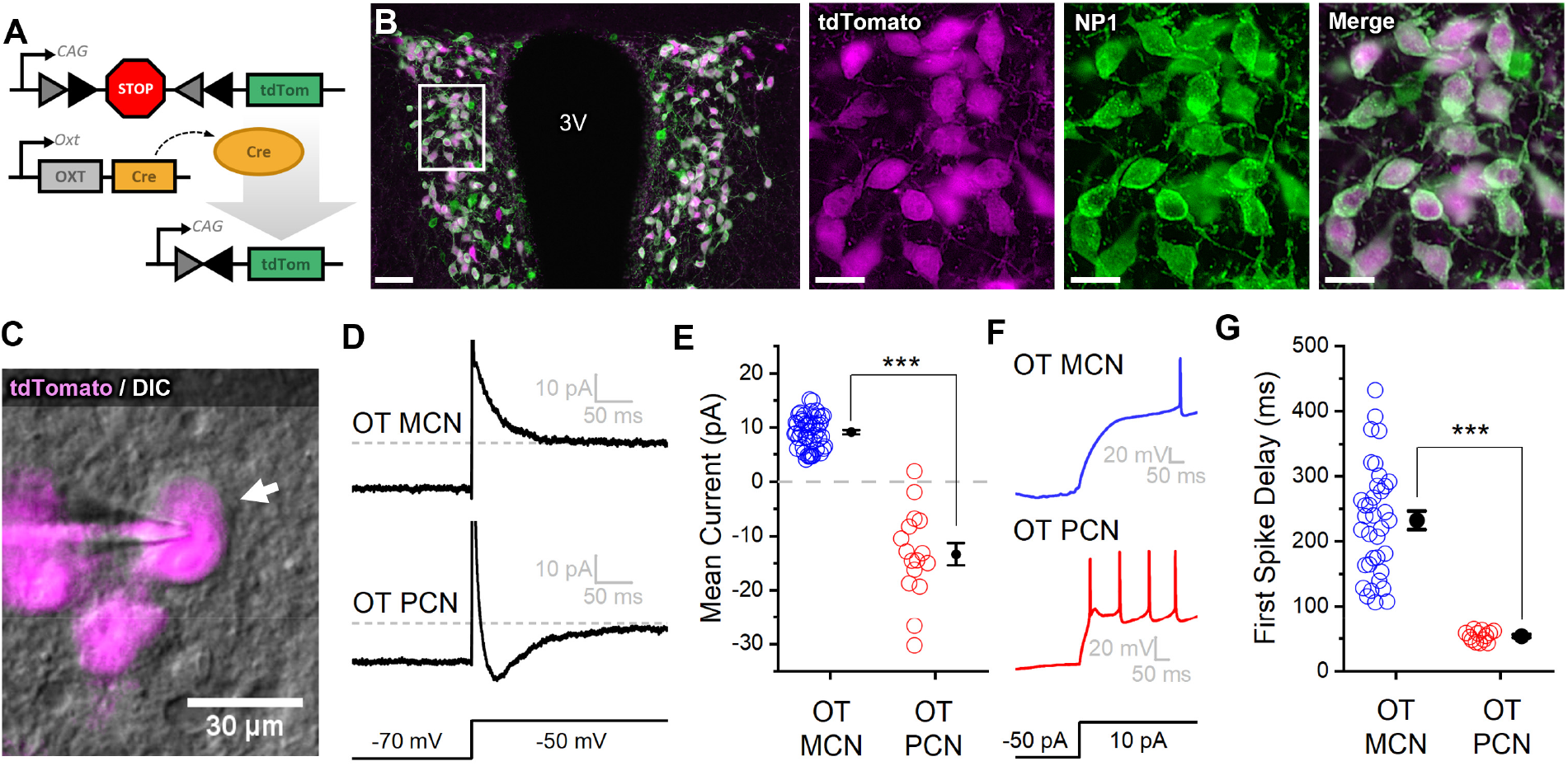
An oxytocin (OT) reporter mouse line facilitates selective targeting of OT neurons for in vitro experiments. **A)** Fluorophore (tdTomato) is selectively expressed in OT neurons of OT-Cre mice using a stop-floxed tdTomato construct. **B)** Immunohistochemistry demonstrates colocalization of tdTomato and NP1 in PVN neurons. See Results for additional details. **C)** Endogenous tdTomato facilitates selective targeting of OT neurons in acute brain slices for whole-cell patch clamp recording. **D)** Magnocellular neurons (OT-MCNs, top panel) can be distinguished from parvocellular neurons (OT-PCNs, bottom panel) by the presence of a transient outward current (*I_A_* current) following a voltage-clamp step from −70 mV to −50 mV. Outward current in OT-MCNs is quantified over the first 100 msec after the step to −50 mV, relative to the steady state current at that voltage (which is indicated by the dashed line. By contrast, OT-PCNs often reveal a transient inward current in response to the same stimulus (quantified with the same strategy, bottom panel). **E)** Mean current response to the voltage step as presented in panel D segregates MCNs (demonstrating a transient outward current, I_A_) from PCNs (with no or inward transient current, n=51, 16, t=10.71, p<0.001). **F)** The *I_A_* current in MCNs produces a delay to first spike which can be readily observed in response to a suprathreshold dep0larizing step in current clamp (top panel). **G)** Time to first spike is significantly longer in OT-MCNs vs. OT-PCNs (n=36, 13, t=12.52, p<0.001). Scale bars: B left panel 100μm, B right panels 20 μm, C 30 μm. Error bars represent mean ± SEM.

### Acute osmotic stress increases basal firing rate of OT-MCNs in vitro

Significant prior evidence suggests that OT-MCNs are likely to be osmosensitive, and further suggest that osmosensitivity may reveal mechanisms that allow OT-MCNs to modulate the relationship between cellular activity and release of OT into the CNS. In order to directly evaluate the osmosensitivity of PVN OT-MCNs in vitro, we used whole-cell current clamp recordings, in combination with bath application of mannitol (MT, an inert osmolyte), to evaluate the effect of acute hyperosmotic stress on firing frequency. We found that bath application of 30 mOsm MT for 5 minutes (in the presence of glutamate and GABA receptor antagonists, see Methods) increased basal firing rate of OT-MCNs by 3.37 ± 0.55 Hz (n = 9, t=6.2, p = 0.0002, Fig. 2A-B, and Extended Data 2-1.) A smaller but longer lasting hyperosmotic stimulus (15 mOsm MT for 10 mins) increased basal firing rate by 1.39 ± 0.55 Hz (n=9, t=2.6, p=0.03, Fig. 2B-C), and was chosen as a standard hyperosmotic stimulus throughout the rest of the study. This effect of MT is cell type specific, as 30 mOsm MT failed to produce a similar effect in PVN OT-PCNs (ΔFreq: 0.7 ± 0.3 Hz, t=2.49, p=0.06 vs. null hypothesis of mean = 0, t=3.58, p=0.005 vs. effect of 30 mOsm MT observed in OT-MCNs, Fig. 2C). Although additional synaptic input from osmosensitive circumventricular organs might further enhance OT-MCN activity in vivo during times of acute osmotic stress, the present results demonstrate PVT OT-MCNs are directly osmosensitive, and further highlight that the primary effect of acute osmotic stress, when observed somatically, is excitatory.

**Figure 2.**
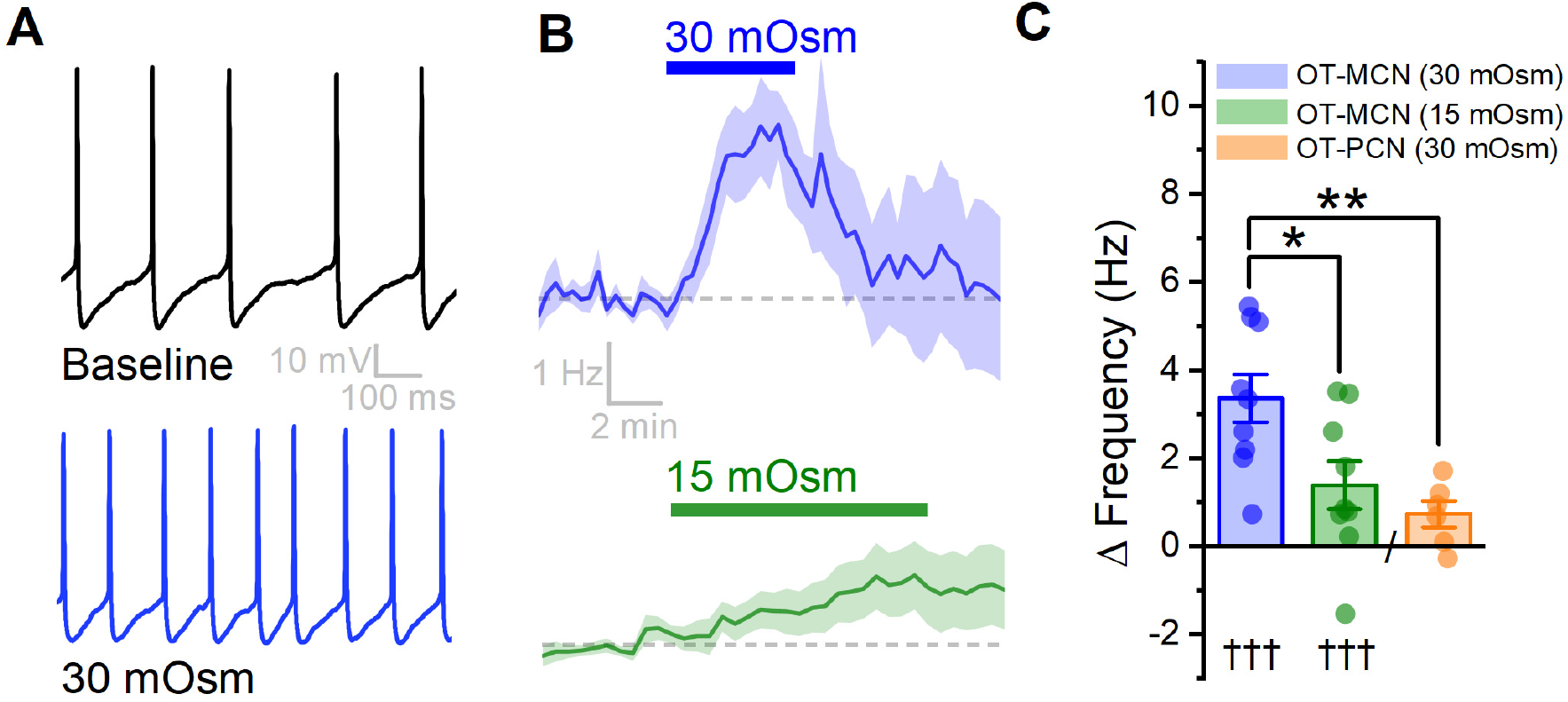
An acute hyperosmotic stimulus increases OT-MCN activity. **A)** Spontaneous firing in representative OT-MCN before (top) and after (bottom) exposure to MT (30 mOsm). **B**) Change in spontaneous action potential frequency over time in response to +30 mOsm (n=9, top) and +15 mOsm (n=9, bottom). **C)** Both 15 and 30 mOsm stimuli caused a significant increase in action potential frequency in OT-MCNs (denoted by †††, p<0.001), but response to 30 mOsm stimulus was significantly larger (denoted by * p< 0.01, blue vs. green bar). By contrast, the same 30 mOsm stimulus did not produce a significant increase in firing rate in OT-PCNs (orange bar, p=0.06 vs. null hypothesis of mean=0), and these results were significantly different that those observed in OT-MCNs (** denotes p = 0.003).

### OT-MCN dendrites are weak conductors of somatic voltage changes

It is clear from prior literature that dendritic release of OT from MCNs is calcium dependent, and it is often presumed that somatic action potentials lead to dendritic calcium influx in a way that promotes dendritic release of peptide (Scala-Guenot, 1987; Fisher and Bourque, 1996b; Tobin et al., 2012a). That said, the relationship between somatic activity and dendritic release of OT is clearly flexible (see Discussion), and key aspects of dendritic physiology potentially impacting this relationship have not been directly examined before. For these reasons, we used a combination of electrophysiological and optical techniques to quantify activity-dependent calcium influx along the length of OT-MCN dendrites as observed in response to a two-second train of action potential like voltage steps delivered to the soma at 20 Hz (Fig. 3A, and Methods). This approach revealed that activity-dependent calcium influx drops rapidly with increasing distance from the soma (F1.6,,6.4=15, p=0.005, Fig. 3B, Fig. 3C, blue trace), suggesting that under basal conditions OT-MCN dendrites lack the ability to actively propagate somatic voltage changes to distal locations, and likely also have a relatively low membrane resistance. However, similar data might be expected if calcium indicators had failed to perfuse into the distal dendrites, or if distal dendrites expressed substantially fewer voltage-gated calcium channels. To evaluate both possibilities simultaneously we repeated the experiment using a cesium gluconate internal solution (see Methods) to block cesium sensitive leak channels and thereby increase dendritic membrane resistance. Notably, in these conditions we observed an overall increase in activity-induced calcium influx (F_2.7,23.9_=9.45, p= 3.9 x 10^-4^ vs. observations in K-glu, Fig. 3C, red trace), with post-hoc tests indicating increased response at distances > 60 μm from the soma). These results clearly indicate that functional calcium indictor is present in the distal dendrites, and that voltage-gated calcium channels are still robustly expressed at distal locations. As such, they also substantially reinforce the conclusion that loss of activity-dependent calcium influx at increasing distance from the soma, as observed when using a more physiological K-gluconate based internal solution, was produced by loss of current through open channels along the dendritic membrane.

**Figure 3.**
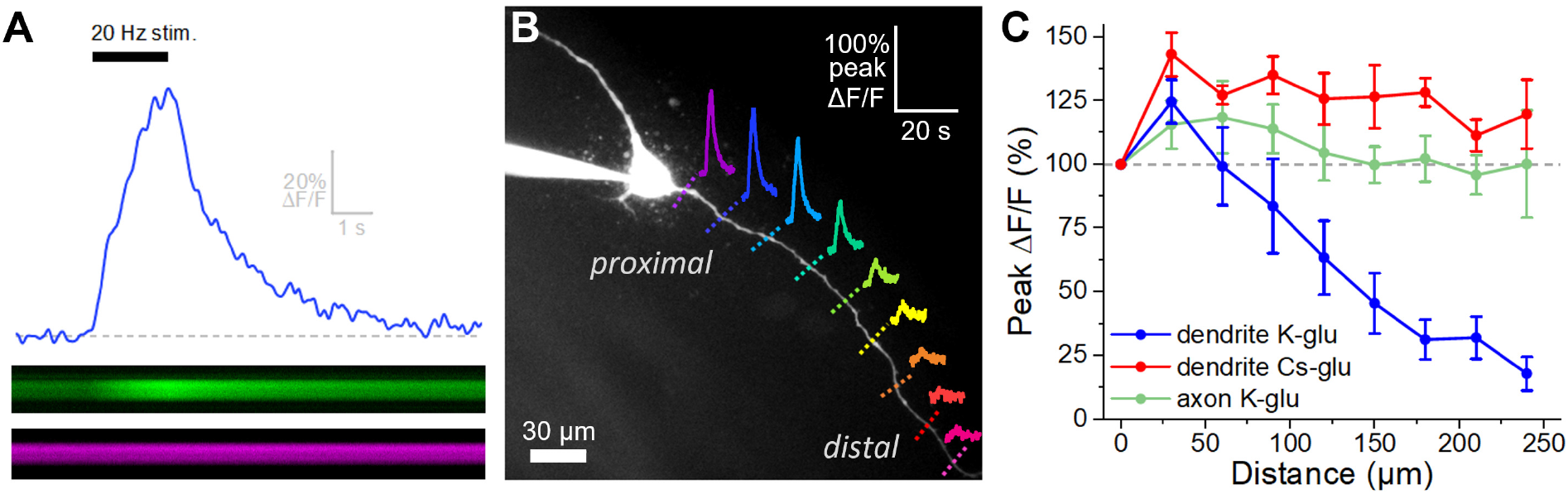
Activity-dependent dendritic calcium influx decreases with distance from the soma in OT-MCNs. **A)** Blue trace illustrates calcium response observed in an OT-MCN dendrite as produced by a 2-second train of action potential-like voltage steps delivered to the soma at 20 Hz (black bar). The calcium signal was calculated using emissions from a calcium sensitive and calcium insensitive indicator (Fluo-5F and Alexa Fluor 594, respectively) obtained by performing a two-photon line scan across the dendrite during somatic stimulation. The bottom panels illustrate line scan data (top: Fluo-5F, bottom: Alexa Flour 594), with time on the horizontal axis and space across the dendrite on the vertical axis. See methods for details. **B)** Two-photon Z-series projection of a representative OT-MCN. Colored dashed lines indicate position of line scans performed in this cell during somatic stimulation as in panel A. Solid lines illustrate calcium response, calculated as in panel A, observed at each location. Scale bar in the top right applies to all traces. **C)** Illustrates the peak of the calcium response as a percentage of baseline (most proximal response) plotted against distance from the soma. Data were obtained from OT-MCN dendrites using either a K-gluconate or Cs-gluconate based internal solution (blue vs. green traces, respectively), and from the axon of OT-MCNs (red trace). Overall, these data reveal significant distance-dependent loss of activity-induced calcium influx only in OT-MCN dendrites, and only in cells that were not filled with a Cs-gluconate based internal solution.

Next, we repeated the experiment for a third time, using a K-glu internal, to evaluate activity-dependent calcium influx along the length of OT-MCN axons. Axons were distinguished from dendrites based primarily on their smaller initial diameter as observed with 2P epifluorescence microscopy (e.g. Fig. 4A). However, consistent with prior reports (Hatton, 1990; Stern and Armstrong, 1998), we also noted that axons often (but not always) branch off a primary dendrite very close to the soma. The results of this experiment indicated that OT-MCN axons, unlike dendrites, reliably and actively propagate somatic voltage changes in cells filled with potassium gluconate (Fig. 3C, green trace). Specifically, a two-way repeated measures ANOVA revealed a main effect of structure on activity-induced calcium influx (axon vs. dendrite, both with K-Glu based internal, F_3.2, 13.1_=9.71, p= 0.001), while post-hoc tests revealed that significantly less loss of response was observed in axons compared to dendrites at distances > 90 μm from the soma. Interestingly, these data also highlight robust expression of voltage-gated calcium channels along the length of OT-MCN axons.

**Figure 4.**
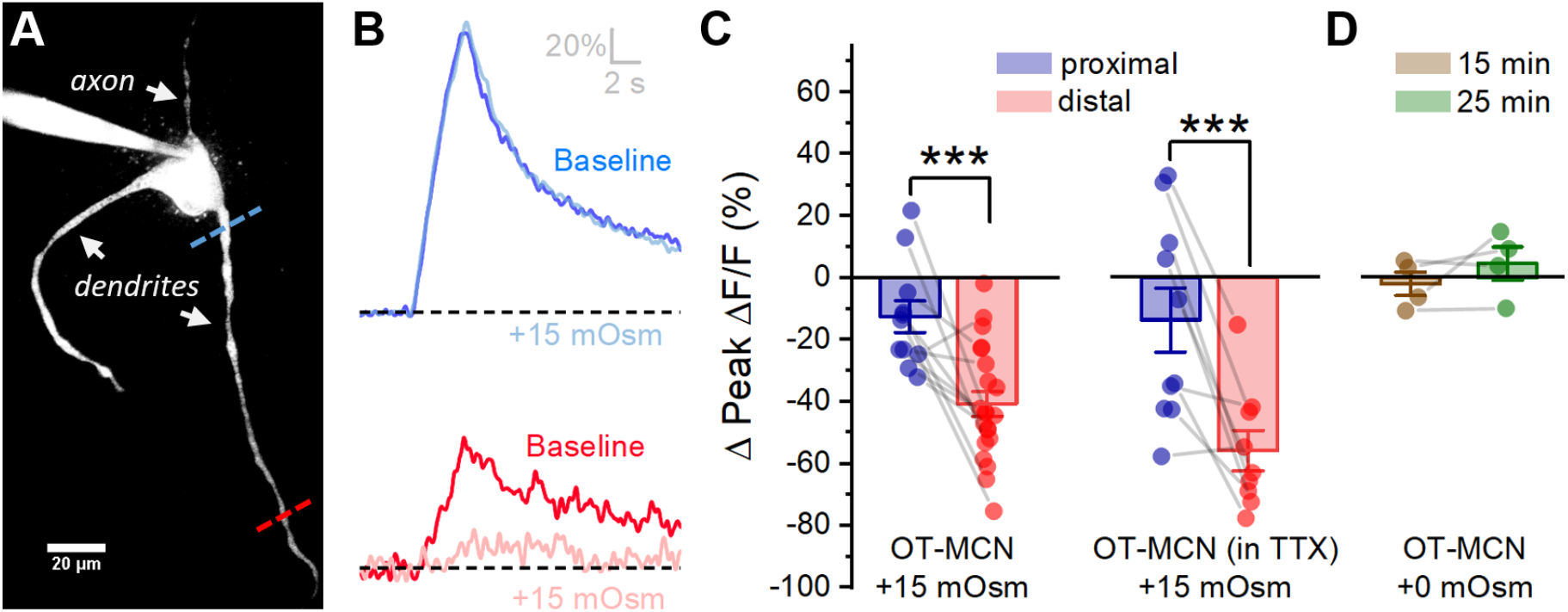
Acute hyperosmotic stress produces a distance-dependent reduction in activity-dependent calcium influx, as observed in OT-MCN dendrites. **A)** A two-photon Z-series projection image of a representative OT-MCN indicating proximal (blue) and distal (red) calcium imaging locations. **B)** Activity-induced calcium response observed before (dark blue) and during (light blue) exposure to a 15 mOsm stimulus at proximal and distal sites indicated in panel A. **C)** Across multiple cells tested, the 15 mOsm stimulus had a significantly greater inhibitory effect on activity-induced calcium influx as observed in the distal vs. proximal dendrites (Left panel, *** p=0.002). Although raw calcium influx was reduced in cells pretreated with 1 μM TTX, a 15 mOsm stimulus continued to preferentially inhibit activity-dependent calcium influx as observed in the distal dendrites (right panel, *** p=0.003). D) Absent hyperosmotic stimulus, activity-dependent calcium influx measured as in panels A-C is stable in both proximal and distal dendrites over a 25 min recording period (sufficient time to change bath conditions in earlier experiments).

### Acute osmotic stress preferentially reduces activity-dependent calcium influx in distal vs. proximal dendrites of OT-MCNs

We next tested the hypothesis that acute hyperosmotic stress (as produced by bath application of 15 mOsm MT) directly modulates the relationship between somatic activity and activity-dependent dendritic calcium influx in OT-MCNs. Towards that end, we used a technical approach very similar to that employed in Fig. 3, however instead of measuring activity-induced influx at multiple locations under control conditions, we picked just two dendritic locations (proximal and distal to the soma) and repeatedly measured activity-induced calcium influx before and after acute hyperosmotic stimulation. Proximal dendritic locations were located within 25 μm of the soma, while distal ones were located at ~125 μm from the soma (See Fig. 4A, blue and red dashed lines, respectively). In order to generate activity-induced calcium influx at these locations the soma was stimulated with the same 2-sec 20 Hz train of action potential like voltage steps as used in Fig. 3, 2P line scan data were collected from each dendritic location and analyzed in an identical manner, and experiments were again performed in the continuous presence of bath applied antagonists for glutamate and GABA receptors (see previous Results section and Methods). We found that acute hyperosmotic stimulation had a mild inhibitory effect on activity-induced calcium influx in the proximal dendrites of OT-MCNs (−12.6 ± 5.15 %, n = 11, t = −2.4, p = 0.034), and yet strikingly, had a much stronger inhibitory effect in the distal dendrites (−40.8 ± 4.0%, n = 21, t = 4.2, p = 0.0002 vs. proximal, Fig. 4B, Fig. 4C, left panel, see Extended data Fig. 4-1 for data separated by sex). Notably these changes were not associated with any significant impact on somatic membrane resistance (baseline: 1,435 ± 187.5 MΩ, after MT: 1,357 ± 177.9 MΩ, n=21, t=0.3, p=0.8), or on the voltage clamp current applied to the soma during the train (94.9 ± 5.21% of baseline after MT, n=11, t=-1.0, p=0.4). Considered together, these observations strongly suggest a site of action along the dendritic membrane. Although glutamate and GABA receptors were antagonized during this experiment, it seemed conceivable that hyperosmotic stress might promote action potential dependent release of other modulators (from cells other than the one patched) which then act locally on OT-MCN dendrites to reduce activity-dependent calcium influx. In order to test this possibility, we repeated the experiment above with 1 μM TTX in the bath (in addition to glutamate and GABA receptor antagonists). However, under these conditions, we found that acute hyperosmotic stress continued to robustly and preferentially inhibit activity-induced calcium influx in the distal vs. proximal dendrites (by −55.8 ± 6.53 vs. −13.7 ± 10.37%, distal, proximal, n=9,10, respectively, t=3.35, p=0.004, Fig. 4C, right panel). Next, in order to eliminate any unexpected impact of the extra time required to change bath conditions in this experiment, we measured activity-dependent calcium influx exclusively in the distal dendrites over 25 minutes under constant bath conditions and noted that there was no significant change (−2.0 ± 3.8 %, 4.6 ± 5.3 %, at 15 and 25 min, respectively, n=4, 4; t=-0.51708, 0.86; p=0.64, 0.45, Fig. 4D). Collectively, these data clearly indicate that despite having an excitatory effect on action potential frequency as observed in the soma (Fig. 2), acute hyperosmotic stress also preferentially reduces activity-dependent calcium influx as observed in the distal vs. proximal dendrites of OT-MCNs, strongly suggesting a change in dendritic membrane resistance.

### Effects of acute osmotic stress on activity-dependent calcium influx in distal dendrites are very likely mediated by changes in dendritic membrane resistance

To further strengthen the conclusion that the results in Fig. 4 are produced by an osmosenstive change in dendritic membrane resistance, we performed an additional control experiment designed to further rule out alternative mechanisms. Specifically, we initiated whole-cell patch clamp recordings from OT-MCNs using the same K-glu internal as in Fig. 4, but now without glutamate receptor antagonists in the bath. We then stimulated them alternately (every 15 seconds) with two distinct stimuli (one delivered to the soma and one to the distal dendrites). The somatic stimulus was the same 2 sec 20 Hz train of action potential like voltage steps used in Fig. 3–4, while the dendritic stimulus was focal application of exogenous glutamate (accomplished using a picospritzer, See Methods). For each stimulus, responses were measured in both the soma (as a whole-cell current) and in a distal dendrite (as ΔG/R generated with a 2P line scan as in Figs. 3–4). Collectively, this experimental design (Fig. 5A) provides insight, in each individual OT-MCN tested, on how acute hyperosmotic stress effects current propagating along the dendrite in both directions, either from the soma towards the distal dendrites, or from the distal dendrites towards the soma. The results emphasize that acute hyperosmotic stress consistently and selectively inhibits whichever response is measured distal from the stimulus that produced it, irrespective of whether that response is measured using electrical or optical techniques. For example, hyperosmotic stress reduced somatic current observed in response to exogenous glutamate delivered to the distal dendrite by 62.5 ± 10.1% (n = 5, t = −6.17, p = 0.003, Fig. 5B), and yet had no effect on somatic current observed in response to a train of AP like voltage pulses delivered to the soma (ΔCurrent = −2.4 ± 4.53%, n = 10, t = −0.5, p = 0.612, Fig. 5B). Conversely, hyperosmotic stress reduced dendritic calcium influx produced by delivering a train of voltage pulses to the soma (by −33.13 ± 7.5 %, n = 10, t = −4.4, p = 0.002, Fig. 5C), and yet had no effect on dendritic calcium influx observed in response to locally applied glutamate (Δ Peak Δ F/F: 1.94 ± 8.7 % of baseline, n = 10, t = 0.22, p = 0.828, Fig. 5C). Overall, we believe that these results, in combination with other data presented above, effectively rule out the idea that the observed effects of acute hyperosmotic stress depend on direct modulation of voltage-gated calcium channels, calcium induced calcium release, or on other aspects of calcium homeostasis in the dendrites. As such, they further strengthen the conclusion that acute osmotic stress is directly modulating dendritic membrane resistance (and thus voltage propagation) in OT-MCNs. Finally, we attempted to further confirm this interpretation using dual path clamp recordings from the soma and distal dendrites of a single OT-MCN. This approach has been used successfully to study the physiology of large dendrites in rat cortical pyramidal neurons and in cerebellar Purkinje cells, however existing literature highlights that dendritic patching in smaller multipolar neurons is only feasible at proximal locations (Davie et al., 2006). To the best of our knowledge, no prior studies have successfully used electrophysiological tools to directly record from the distal (or proximal) dendrites of PVN neurons. Indeed, we found that the very small diameter and variable orientation of mouse OT-MCN distal dendrites, combined with tissue movement associated with changes in osmotic pressure, prohibited direct measurement of dendritic membrane resistance in this manner. Thus, we conclude that the combined electrical / optical approaches used in this study effectively and reliably reveal novel aspects of dendritic physiology in OT-MCNs that are not accessible to direct electrical recording.

**Figure 5.**
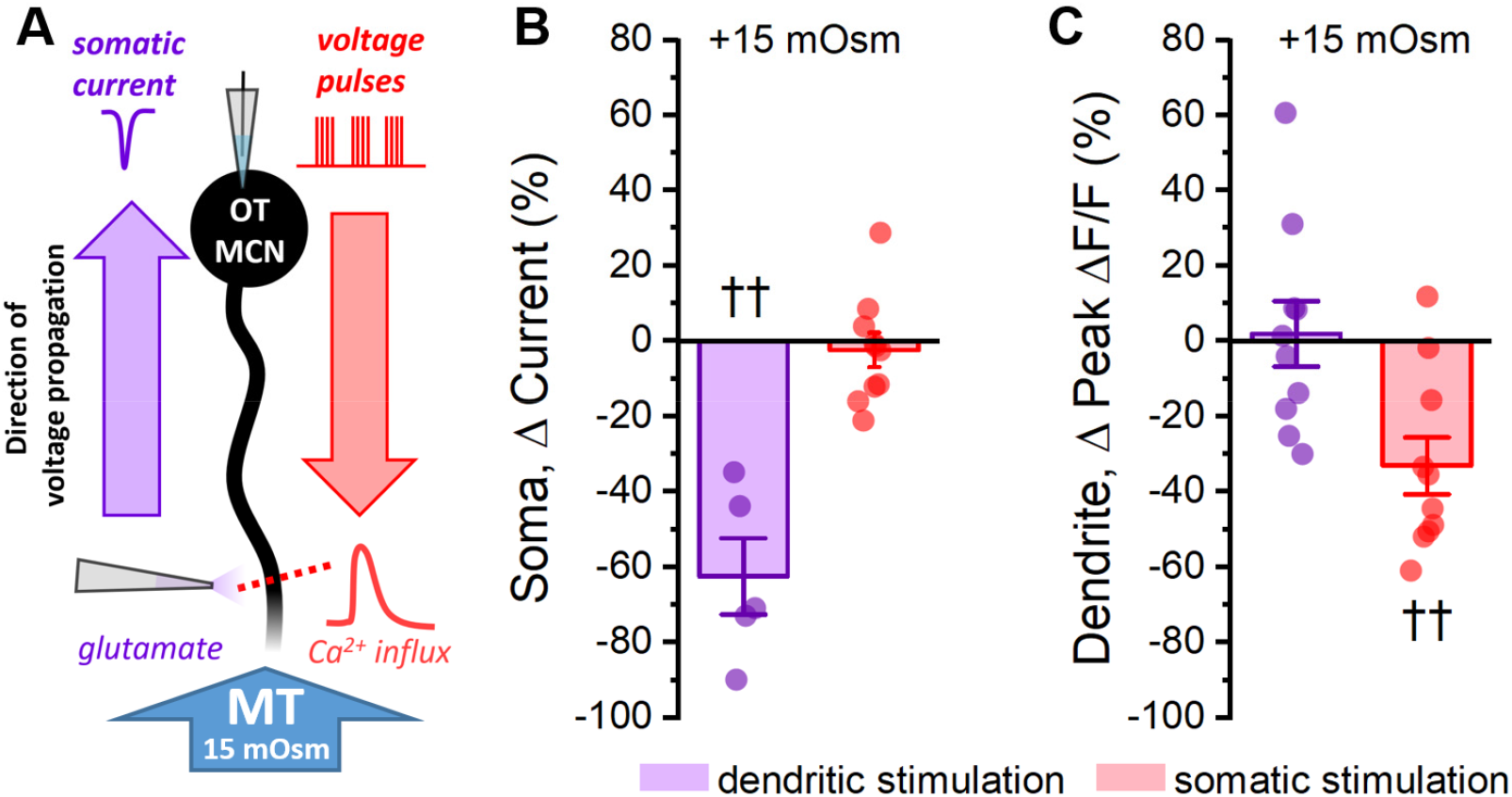
Acute hyperosmotic stress selectively inhibits responses measured distal from a stimulus. **A)** Diagram of experimental design. Two distinct stimuli (one to the distal dendrites, and one to the soma) were delivered every 15 seconds to an OT-MCN voltage clamped at −70 mV. The stimulus to the distal dendrites was local application of glutamate (bottom left), while the somatic stimulus was the same 2-sec, 20 Hz train of action potential-like voltage pulses used in earlier experiments (delivered though the patch pipette, top right). For each stimulus, we used electrophysiological techniques (whole-cell recording) to measure the somatic response, and optical techniques (two-photon line scans across the distal dendrite as in earlier figures) to measure the dendritic response. Both the electrophysiological and optical responses to each stimulus were measured both before and after bath application of 15 mOsm MT. **B)** Illustrates the effect of acute hyperosmotic stress on somatic responses to either dendritic or somatic stimulation (purple, red, respectively). **C)** Illustrates the effect of acute hyperosmotic stress on dendritic responses to either dendritic or somatic stimulation (purple, red, respectively). Collectively these results highlight that bath application of 15 mOsm MT effectively and selectively inhibits responses that are measured distal from the stimulus that produced them, irrespective of whether those responses are measured with electrical or optical techniques. These data reinforce the conclusion that acute hyperosmotic stress reduces dendritic membrane resistance. †† p<0.01.

### Changes in dendritic membrane resistance produced by acute hyperosmotic stress are cell type specific, compartment specific, and bidirectional

Next, to eliminate any possible generalized or nonspecific effects of acute hyperosmotic stress, we designed experiments to test the hypothesis that the effects are both compartment specific and cell type specific. Compartment specificity was evaluated using techniques identical to those employed for Fig. 4, except that we compared activity-dependent calcium influx in the distal dendrite to that observed in the distal axon. We found that acute hyperosmotic stress again effectively inhibited activity-dependent calcium influx in distal dendrites (Δ Peak Δ F/F: −44.0 ± 8.0 %, n = 5, t = −5.5, p = 0.005), and yet produced a much smaller effect in the distal axon (Δ Peak Δ F/F:-10.0 ± 2.3 %, n = 5, t = 3.4, p = 0.027 vs. distal dendrite, Fig. 6A). This result demonstrates significant compartment specificity within individual OT-MCNs. In order to evaluate cell type specificity, we used identical approaches to measure activity-dependent calcium influx in proximal vs. distal dendrites in PVN OT PCNs (identified as described in Fig. 1), and in CA1 pyramidal cells. In OT PCNs there was no significant effect of acute hyperosmotic stress in proximal or distal dendrites (Δ Peak Δ F/F: −7.7 ± 4.3 %, n = 5, t = −1.8, p = 0.147; 6.6 ± 3.4 %, n = 5, t=1.96, p=0.12, respectively, Fig. 6B). In CA1 pyramidal cells, we noted a mild inhibitory effect in proximal dendrites, but no effect in distal dendrites (−6.4 ± 2.4 %, t = −2.7, p = 0.045; −3.80 ± 3.5 %, n = 6, t = −1.1, p = 0.328, respectively, Fig. 6C). These results effectively demonstrate cell type specificity. Next we reasoned that if acute hyperosmotic stress inhibits calcium induced influx in the distal dendrites of OT-MCNs by opening a distinct dendritic osmosensitive ion channel, and if that channel is not completely closed in control conditions, then an acute hypoosmotic stimulus should have opposite effects. Indeed, we found that acute reduction of osmolarity by 30 mOsm (achieved by diluting the bath solution with water) effectively increased activity-dependent calcium influx in the distal dendrites of OT-MCNs (by 31 ± 10.7 % of baseline, n = 10, t = 3.0, p = 0.014, Fig. 6D), while having no effect in the proximal dendrites (n = 10, t = −0.6, p = 0.536, Fig. 6D). As with effects of hyperosmotic stress, these changes occurred absent any significant effect on somatic membrane resistance or voltage clamp current observed during stimulation (n=10, t=1.3, p=0.21; n=10, t=-1.84, p=0.1, respectively). Collectively, these results demonstrate that the effects of acute osmotic stress on activity-induced calcium influx, as observed in the distal dendrites of OT-MCNs, are compartment specific, cell type specific, and bidirectional.

**Figure 6.**
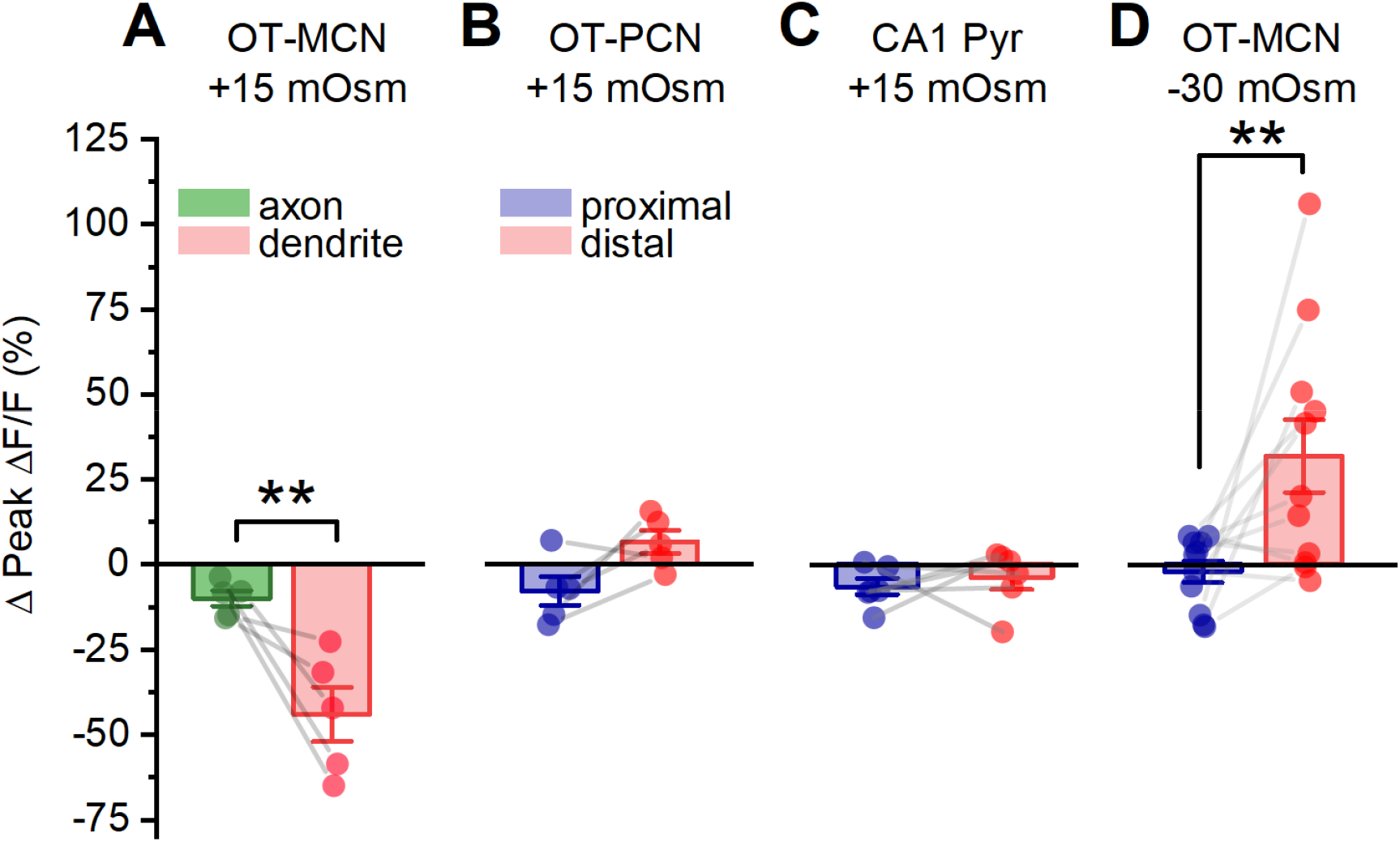
Effect of acute hyperosmotic stress is compartment-specific, cell type-specific and bidirectional. **A)** Compartment specificity: Acute hyperosmotic stress preferentially inhibits activity-dependent calcium influx in the distal dendrites as opposed to the distal axon of OT-MCNs. **B-C)** Cell type specificity: In contrast to results reported for OT-MCNs (Fig. 4), acute hyperosmotic stress does not preferentially or selectively inhibit calcium influx (observed using identical techniques) in the distal dendrites of either PVN OT-PCNs (panel B) or hippocampal CA1 pyramidal neurons (panel C). **D)** Bidirectionality: In contrast to results reported in Fig. 4, acute hypo- (rather than hyper-) osmotic stress selectively and preferentially increases (rather than decreases) activity-dependent calcium influx observed in the distal dendrites of OT-MCNs. As in prior experiments on distal dendrites, all line scans for these experiments were conducted at ~125 μm from the soma, and acute hyperosmotic stress was produced by bath application of 15 mOsm. Acute hypoosmotic stress (of −30 mOsm) was produced by diluting ACSF in the bath with water. ** p≤0.01.

### Changes in dendritic membrane resistance as produced by acute osmotic stress preferentially inhibit minimally evoked EPSCs produced by a distal vs. proximal stimulator

Data presented in Fig. 5 indicates that the somatic response to activation of glutamate receptors on the distal dendrites of OT-MCNs is significantly inhibited by acute osmotic stress. This result (in combination with other results above) suggests that changes in dendritic membrane resistance are likely to impact integration of synaptic inputs, as well as activity-dependent calcium influx. In order to test this hypothesis directly we used minimal stimulation techniques (See Methods) to evoke glutamate release from one or few axons that make synaptic contact with either the proximal or distal dendrites of an OT-MCN (Fig. 7A). After identifying a clear evoked excitatory postsynaptic current (eEPSC), we bath applied 15 mM MT as in prior experiments. We found that acute hyperosmotic stress reliably and reversibly reduced eEPSC amplitude as evoked by a minimal stimulator placed near the distal dendrite (by 63.4 ± 8.7%, n = 8, t = −7.3, p = 1.6 x 10^-4^, Fig. 7A,C). As in earlier experiments, this result was not associated with a significant change in somatic membrane resistance (n=8, t=-0.14, p=0.90). If, as expected, the effect is instead produced primarily by a drop in dendritic membrane resistance, then the same acute osmotic stimulus should have less of an inhibitory effect on EPSCs generated using a minimal stimulator placed near the proximal dendrite. Indeed, consistent with this hypothesis, we found that 15 mM MT reduced proximally evoked EPSC amplitude by 30 ± 9.6% (n=6, t=-3.16, p=0.03, Fig. 7A,B). Consistent with our hypothesis, this effect is significantly smaller than observed when using a stimulator placed near the distal dendrite (t=-2.55, p=0.03). In order to confirm that eEPSCs involved in these experiments were glutamatergic, a subset of MT sensitive responses (n=3) were challenged with bath applied glutamate receptor antagonists after recovery, and were effectively eliminated (not illustrated).

**Figure 7.**
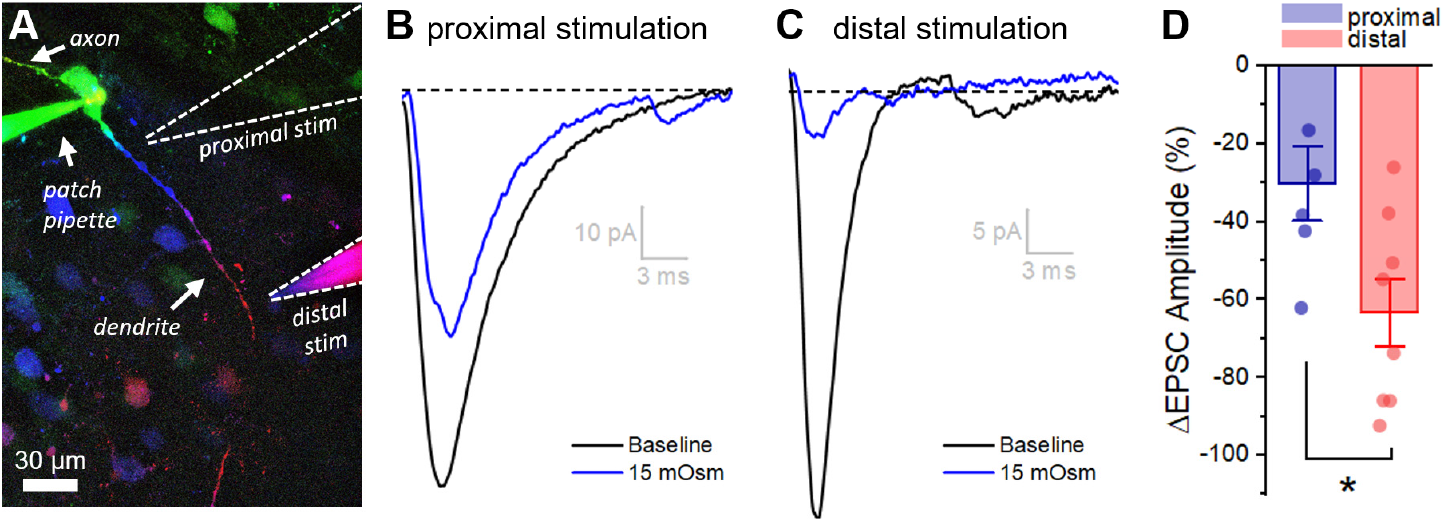
Acute hyperosmotic stress has a greater inhibitory effect on distally vs. proximally evoked EPSCs in OT-MCNs. **A)** Representative OT-MCN (depth-coded two-photon Z-series projection) illustrates placement of minimal stimulators used to evoke EPSCs at either proximal or distal dendritic locations. **B-C)** Representative data illustrating evoked EPSCs produced by proximal (panel B) vs. distal stimulation (panel C), both before and after bath application of 15 mOsm MT. **D)** Summary data indicates that on average, acute hyperosmotic stress produces greater inhibition of distally evoked EPSCs. * p=0.03.

### Distinct somatic vs. dendritic effects of acute osmotic stress on OT-MCNs are likely to depend on compartment specific and molecularly distinct osmosensors

Results presented in figures 3–6 suggest that the effects of acute hyperosmotic stress on activity-dependent calcium influx in distal dendrites of OT-MCNs are likely mediated by an osmosensitive ion channel expressed along the dendritic membrane. By contrast, results presented in Fig. 2, as well as significant prior literature more broadly focused on osmosensation in MCNs, suggest that a functional somatic osmosensor likely also exists (Bourque, 2008). Based on the nature of the effects observed in the soma compared to the dendrites, we hypothesized that somatic and dendritic osmosensors in OT-MCNs are likely to be molecularly distinct. In order to test this hypothesis directly we evaluated the effect of ruthenium red (RR), a generic antagonist of transient receptor potential cation channels in subfamily V (TRPV receptors), on both somatic and dendritic effects of acute hyperosmotic stress observed in OT-MCNs (as in Figs. 2 and 7, respectively). Consistent with our hypothesis, we found that pre-treatment with 10 μM RR blocked the effect of hyperosmotic stress on basal firing rate as observed in current clamp (Δ action potential frequency: 0.28 ± 0.21, n=7, t=0.5, p=0.61 vs. null hypothesis of mean=0, t=4.0, p=0.001 vs. response to same stimulus absent RR, Fig. 8A-B). Conversely, RR did not block the inhibitory effect of acute hyperosmotic stress on evoked EPSCs as observed in a separate group of cells in voltage clamp (Δ EPSC amplitude: −71.5 ± 6.0%, n=6, t=-11.9, p<0.001 vs. null hypothesis of mean=0, t=0.72, p=0.49 vs. effect observed absent RR, Fig. 8C-D). These results reinforce the hypothesis that PVN OT-MCNs express distinct osmosensors in somatic vs. dendritic compartments, and further suggest that the somatic but not dendritic osmosensor may be a member of the TRPV family. Interestingly, we also noted that the effect of MT on activity-induced calcium influx as observed in the distal dendrites of OT-MCNs is blocked in cells filled with a cesium-gluconate internal solution, which suggests that the dendritic osmosensor may be cesium sensitive (See Extended Data Fig. 8-1 for additional details).

**Figure 8.**
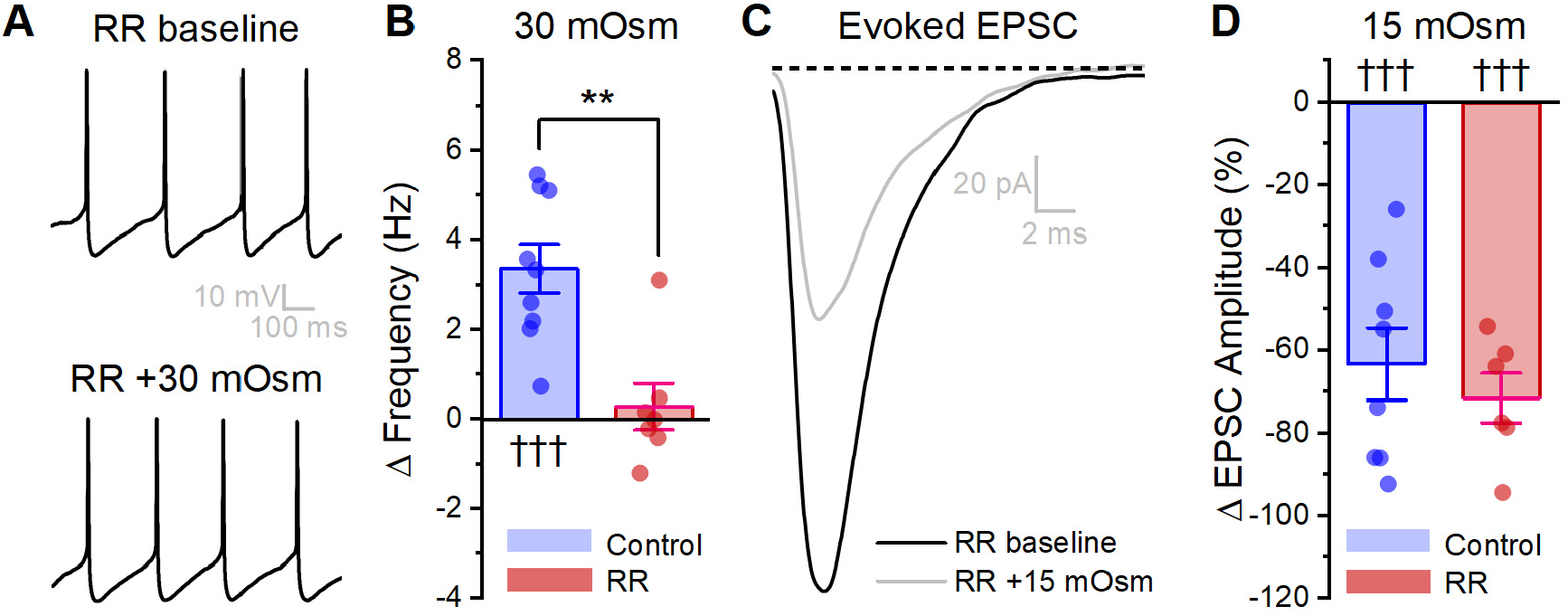
Ruthenium red selectivity blocks the somatic (but not dendritic) effect of acute hyperosmotic stress. **A)** Raw data from a representative OT-MCN illustrating spontaneous firing before and after acute hyperosmotic stress in the continuous presence of TRPV antagonist ruthenium red (RR). **B)** Summary data indicating that the effect of acute hyperosmotic stress on action potential firing frequency in OT-MCNs is blocked in cells pretreated with RR. **C)** Raw data from a representative OT-MCN illustrating that acute hyperosmotic stress continues to inhibit minimally evoked EPSCs evoked at the distal dendrite, even in cells pretreated with RR. **D)** Summary data highlighting that pretreatment with RR has no impact on the ability of acute hyperosmotic stress to inhibit distally evoked EPSCs. Note that control datasets in panels B and D were previously presented in Fig. 2C and Fig. 7D, respectively. ** p=0.001, ††† p≤0.001.

### GABA_A_ receptor activation preferentially reduces activity-dependent calcium influx in distal vs. proximal dendrites of OT-MCNs

Next, we sought to determine whether dendritic membrane resistance in OT-MCNs is an aspect of dendritic physiology that can be actively manipulated to modify the relationship between somatic activity and dendritic calcium influx, even under conditions that do not involve changes in osmotic stress. Towards that end, we performed experiments similar to those presented in Fig. 4, except we replaced the hyperosmotic stimulus with bath application of 400 nM muscimol (a GABA_A_ receptor agonist). Unlike hyperosmotic stress, bath application of muscimol significantly reduced somatic membrane resistance (from 1178 ± 166.6 MΩ to 226 ± 43.5 MΩ, n = 6, t = 4.9, p = 0.004) and produced a tonic inhibitory current (of 15.4 ± 5.42 pA, n = 6, t = 2.8, p = 0.036) apparent in cells voltage clamped at −70 mV, consistent with tonic activation of somatic GABA_A_ receptors. However, importantly, like acute hyperosmotic stress, muscimol preferentially inhibited activity-induced calcium influx in the distal vs. proximal dendrites (−75.8 ± 2.7 % vs. −31.6 ± 6.4 %, respectively, n=6, t=5.6, p=0.003, Fig. 9). This finding is consistent with opening of GABA receptors along the dendritic membrane (see also Pirker et al., 2000; Park et al., 2006), and notably, highlights that dendritic membrane resistance is likely to be under constant and dynamic regulation in OT-MCNs even absent changes in osmotic stress.

**Figure 9.**
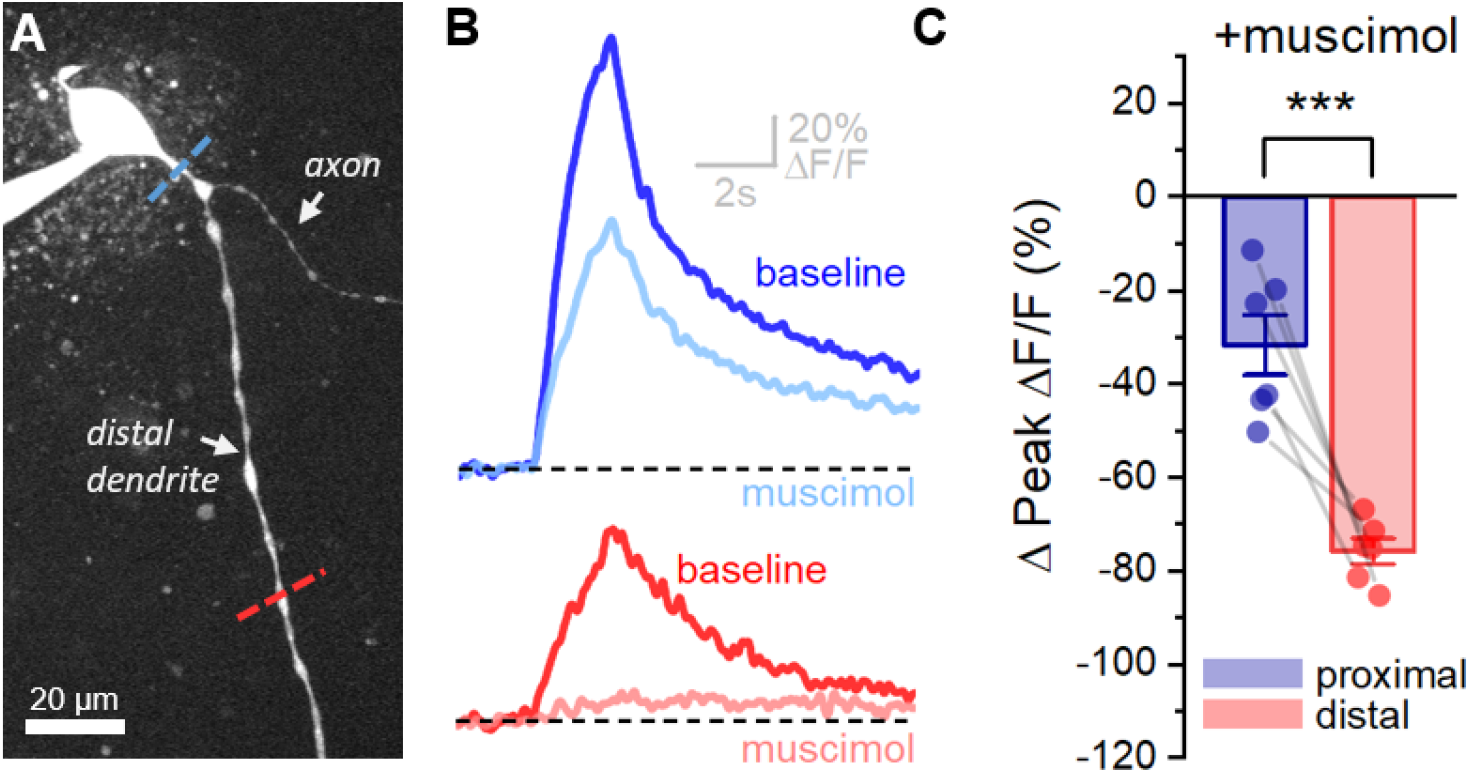
Activation of GABA_A_ receptors preferentially inhibits activity-dependent calcium influx in distal vs. proximal dendrites of OT-MCNs. **A)** Calcium responses to somatic activity were measured at proximal and distal sites (dashed lines) before and after bath application of 300 nM muscimol (a GABA_A_ receptor agonist). Other than using muscimol in place of acute hyperosmotic stress, techniques are identical those described for Fig. 4. **B)** Representative calcium responses observed during somatic stimulation in the proximal (blue) vs. distal (red) dendrite of an OT-MCN, before (lighter trac) and after (darker trace) bath application of muscimol. **C)** Summary data indicates that bath application of muscimol produces greater inhibition of activity-dependent calcium influx in the distal vs. proximal dendrites of OT-MCNs. *** p<0.001.

## DISCUSSION

This study uses a combination of electrical and optical recording techniques to examine activity-dependent calcium influx in the dendrites of PVN OT-MCNs in OT-tdTomato reporter mice. Somatic activity was induced with a well-controlled train of action potential like stimuli delivered to the soma, while activity-dependent calcium influx was measured in OT-MCN dendrites using quantitative 2-photon microscopy. We demonstrate that PVN OT-MCN dendrites are weak conductors of somatic voltage changes in basal conditions in both male and virgin female mice, and importantly, we also find that activity-induced calcium influx in the dendrites is subject to robust modulation by a diverse set of stimuli acting on distinct types of ionophores in the dendritic membrane.

The primary stimulus used in the current study is an acute increase in osmotic pressure. Hyperosmotic stimuli are well-recognized for an ability to increase activity of hypothalamic OT-MCNs, and in so doing, for promoting significant increases in both peripheral and central OT concentration. Increases in peripheral OT concentration are believed to depend on axonal release in the neurohypophysis (which delivers OT directly into the vasculature), while increases in central concentration are likely to depend heavily on dendritic release into the extracellular space, the subarachnoid space, and/or the third ventricle (Bourque, 1991; Ludwig and Leng, 2006; Veening et al., 2010). A curious feature of the neurohormonal response to osmotic stress is that increases in peripheral OT concentration are both rapid and frequency dependent (with concentration closely tracking MCN activity), while increases in central OT concentration are temporally separated from peak MCN firing, often by over an hour (Ludwig et al., 1994; Ludwig, 1998). This is somewhat counter intuitive because like axon terminals, OT-MCN dendrites also contain many large dense core vesicles loaded with oxytocin, and these dendritic vesicles are also subject to both activity and calcium dependent release (Mason et al., 1986; Pow and Morris, 1989; Ludwig et al., 1995; Wang et al., 1995). Notably, other types of stimuli can drive more synchronous release of OT from both axons and dendrites, or can preferentially promote dendritic release (Neumann et al., 1993b; Sabatier et al., 2003). Collectively these types of data effectively highlight that although at least loosely coupled, the relationship between somatic activity and dendritic release of peptide in hypothalamic MCNs is highly flexible.

The most well-established basis for understanding this flexibility invokes a model of conditional priming, whereby specific endogenous modulators, acting directly on the dendrites, are able to prime dendritic vesicles to be more available for activity-dependent release (Morris and Ludwig, 2004; Ludwig and Leng, 2006). However, in the specific case of osmotic stress, maximizing calcium dependent dendritic priming prior to hyperosmotic stimulation was found to increase the amount of dendritic release of OT, remarkably, without altering its time course (Ludwig et al., 2002). This striking result suggests that there must be some aspect of OT-MCN physiology that is activated by hyperosmotic stimulation, that is separate from conditional priming, and that is capable of rapidly yet transiently reducing the probability of activity-dependent exocytosis of dendritic OT.

In that regard, a key finding of the current study is that an acute hyperosmotic stimulus delivered in vitro decreases activity-induced calcium influx in OT-MCN dendrites, while an acute hypoosmotic stimulus increases it. In each case, we noted minimal effect of changing bath conditions on somatic input resistance, or on somatic current observed during the stimulus, and yet changes in osmotic stress had a more robust effect in distal vs. proximal dendrites. We noted no similar inhibitory effect of acute hyperosmotic stimuli on activity-induced calcium influx in the distal axons of OT-MCNs, indicating compartment specificity, or in the distal dendrites of OT-PCNs or CA1 pyramidal cells, demonstrating cell-type specificity. Further, in experiments that involved both somatic and dendritic stimulation techniques, we demonstrated that hyperosmotic stimulation selectively inhibits responses measured distal from the stimulus, irrespective of whether those measurements are made using electrical or optical techniques. Collectively, these data indicate for the first time that osmotic stress modulates activity-dependent calcium influx in OT-MCN dendrites by acting on osmosensitive ion channels expressed along the dendritic membrane. Based on these data, we believe it is reasonable to postulate that this mechanism may reduce activity-dependent dendritic release of OT during times of high somatic activity as induced by acute hyperosmotic stress. As such, it may be interesting for future studies to evaluate whether low levels of dendritic calcium influx produced by somatic activity when dendritic osmosensitive channels are open, or concurrent action of other dendritic modulators, helps promote priming of dendritic vesicles in a way that contributes to enhanced dendritic release once basal osmolarity is restored.

Another aspect of this study worth specifically highlighting is the novel implication that OT-MCNs express molecularly distinct osmosensitive channels in their soma vs. dendrites. This conclusion is supported by the observation that a generic TRPV receptor antagonist blocked the effect of hyperosmotic stress on action potential firing frequency without altering effects on EPSCs produced by a minimal stimulator placed near the distal dendrite. It is further strengthened by the observation that intracellular cesium blocked the effects of hyperosmotic stress on activity-dependent calcium influx in the distal dendrites even through TRPV receptors are cesium permeant (Caterina et al., 1997; Puopolo et al., 2013). Based on these results we hypothesize that the somatic osmosensor is a member of the TRPV family, while the dendritic osmosensor is a cesium sensitive potassium channel. While some prior literature is generally consistent with these hypotheses (Liu et al., 2005; Sharif Naeini et al., 2006; Zhang et al., 2009; Prager-Khoutorsky and Bourque, 2015), very few studies have examined these questions with respect to OT-MCNs in particular.

Next, it is interesting to highlight that all novel aspects of OT-MCN dendritic physiology revealed here, as well as most other core intrinsic features of OT-MCNs observed, were identical in male and virgin female mice, suggesting that they have a fundamental and sex independent role in regulation of oxytocinergic signaling. That said, it seems plausible that aspects of OT-MCN physiology likely relevant to central OT signaling, such as the dendritic membrane resistance, could be regulated in a context and sex specific way. As such, it may be interesting to determine whether activity-dependent calcium influx in the distal dendrites of OT-MCNs is naturally increased in pregnant or lactating females. Indeed, concurrent increases in both peripheral and central release of OT have been reported in response to suckling (Neumann et al., 1993).

Other aspects of this study make two additional important points. First, we demonstrate directly that the effects of acute hyperosmotic stimulation on OT-MCN dendrites are not limited to modulation of activity-dependent calcium influx, but also robustly inhibit the somatic response to endogenous synaptic inputs arriving at the distal dendrites. This finding, in combination with other observed effects, suggests that acute systemic osmotic stress not only transiently reduces the probability of activity-dependent dendritic release of OT into the CNS, but also simultaneously reduces the impact of descending central inputs forming dendritic synapses on MCN firing rate. These changes are expected to effectively but transiently prioritize activity-dependent increases in peripheral OT concentration. Second, we report that an ability to modulate activity-dependent calcium influx by acting on dendritic ionophores is not unique to osmotic stress. Specifically, we note that bath application of a GABA_A_ receptor agonist not only decreases somatic membrane resistance in OT-MCNs but also preferentially inhibits activity-induced calcium influx in the distal vs. proximal dendrites. The later finding is consistent with prior reports that GABAergic receptors are expressed on MCN dendrites (Pirker et al., 2000; Park et al., 2006), and it is important in the context of this study because it highlights that endogenous regulation of dendritic conductivity may represent an important mechanism for regulating central OT concentration even in situations that have nothing to do with osmotic stress. Therefore, we expect that it will be important for future studies to evaluate the ability of additional endogenous modulators to impact conductivity of OT-MCN dendrites.

Finally, as noted in more detail in the Introduction, central OT signaling systems are a promising therapeutic target for a variety of conditions impacting mental health, and yet the best available current strategy for therapeutically modulating activation of central OTRs in humans involves intranasal delivery of an exogenous agonist that has low permeability to the blood brain barrier (Evans et al., 2014; Leng and Ludwig, 2016; Quintana and Woolley, 2016; Quintana et al., 2018). In our view, a better understanding of OT-MCN physiology, particularly as it relates to release of endogenous OT in the CNS, may ultimately lead to improved therapeutic options that are better able to mimic natural concentration, kinetic, and context dependent aspects of central OT signaling. In the current study, measures of activity-dependent calcium influx in OT-MCN dendrites reveals a novel aspect of dendritic physiology likely to underlie the flexible relationship between somatic activity and dendritic release of OT. Future studies may develop new methods for spatially precise quantification of dendritic exocytosis and/or for detection of quantal amounts of OT release very close to the dendritic membrane, which would in turn promote a more direct evaluation of the relationship between dendritic calcium influx and dendritic exocytosis.

## ACKNOWLEDGEMENTS

This work was supported by R01 MH104641 and by R01 HL122494. We thank Christopher Ford for helpful comments on the manuscript.

## FIGURES

**Extended Data Figure 1-1:**
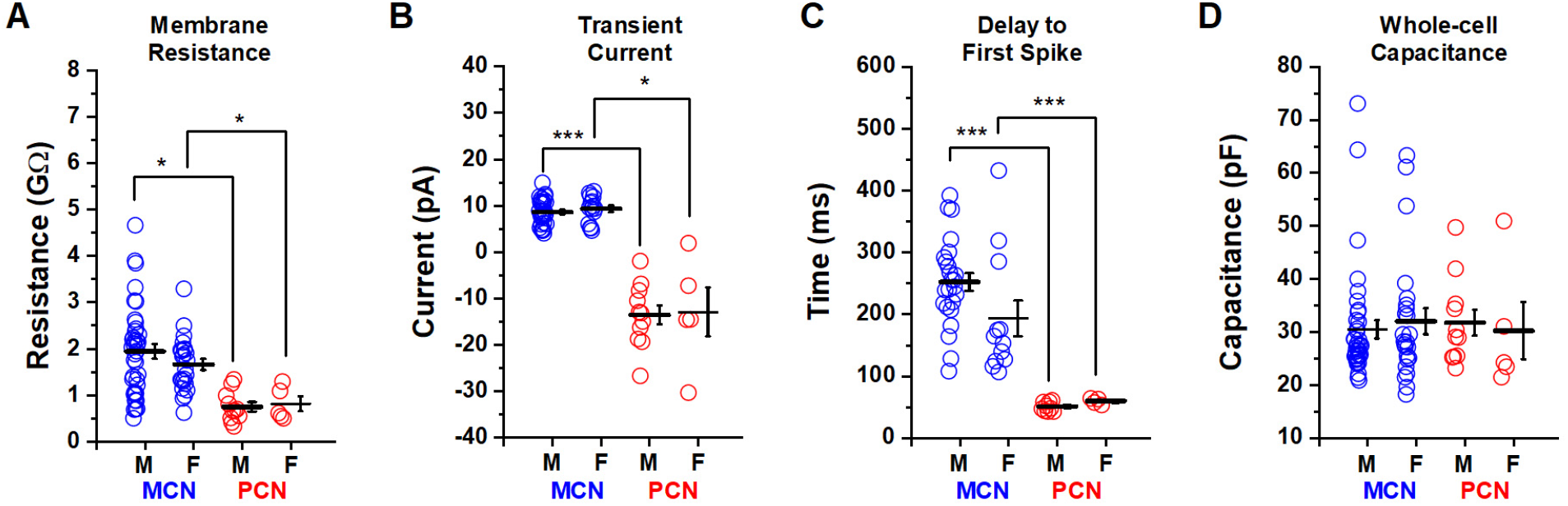
Intrinsic properties of OT PVN neurons do not differ by sex. Core intrinsic properties of OT-MCNs (blue) and OT-PCNs (red) did not differ by sex as illustrated in panels A-D. Although intrinsic properties did not vary by sex within either cell type, 3 of 4 of these properties did differ significantly across cell type. Specifically, in OT-MCNs vs. OT-PCNS, membrane resistance was significantly larger, transient current observed in voltage clamp after a step from −70 mV to −50 mV was outward rather than inward, and delay to first spike observed during a suprathreshold stimulation in current clamp was significantly longer. All three of these differences were apparent in both males and females. Key comparisons between cell types, and within sex, are highlighted by brackets in panels A-C (* = p<0.05, while *** = p<0.001). Somewhat surprisingly, we found that whole-cell capacitance was not significantly different in OT-MCNs vs. OT-PCNs, in either sex (panel D).

**Extended Data Fig. 2-1:**
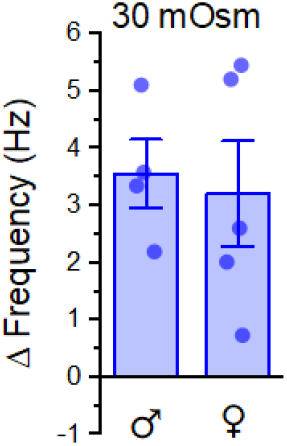
Somatic response to hyperosmotic stress is not different in males vs. females. Mean change in action potential frequency observed in OT-MCNs in response to 30 mOsm MT did not differ between males and females (Male: 3.6 ± 0.6 Hz, female: 3.2 ± 0.9 Hz, n=4,5, t=0.30, p=0.8). Data illustrated here are the same as in blue bar from Fig. 2C, except now separated by sex.

**Extended Data Fig. 4-1:**
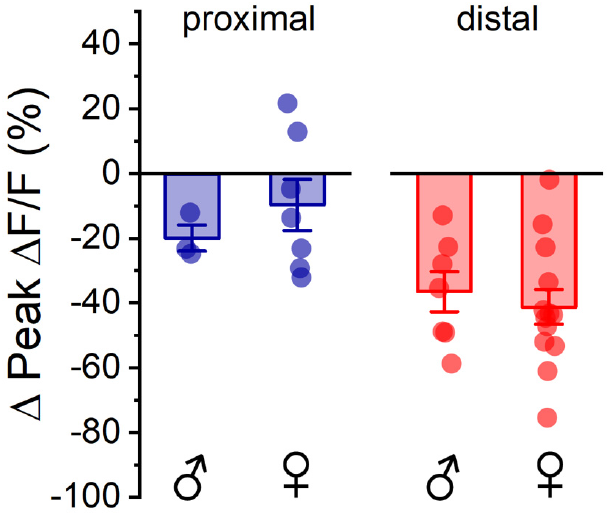
Effect of acute hyperosmotic stress on activity-dependent calcium influx is similar in males and females. The effect of acute hyperosmotic stress on activity-dependent calcium influx, as observed in either proximal or distal dendrites is not sex dependent (proximal: n=3, 7, M,F, respectively, t=-1.2, p = 0.3, distal: n=7, 13, M,F, respectively, t=0.56, p=0.6). As such, these data were combined across sex and presented in Fig. 4C, left panel.

**Extended Data Figure 8-1:**
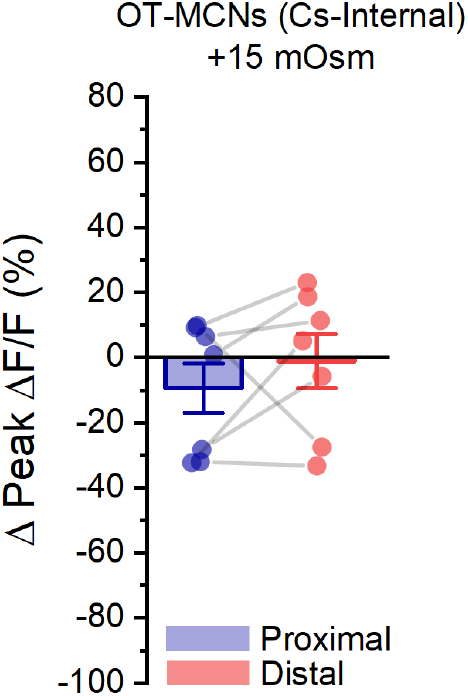
Acute hyperosmotic stress has no effect on activity-dependent calcium influx observed in dendrites of OT-MCNs filled with a Cs-gluconate based internal solution. We noted that when the experiment presented in Fig. 4A-C is conducted in OT-MCNs filled with a Cs-gluconate rather than a K-gluconate based internal solution, acute hyperosmotic stress no longer had any significant effect on activity-dependent calcium influx as observed in either the proximal or distal dendrites (Proximal:. −9.5 ± 7.7%, Distal: −1.2 ± 8.3%, n=7 is both cases, t=-1.2, −0.1, p=0.26, 0.9, respectively, using one-sample t-test against null hypothesis mean=0, and group means were also not significantly different, t=-0.93, p=0.39, paired t-test).

